# spARC recovers human glioma spatial signaling networks with graph filtering

**DOI:** 10.1101/2022.08.24.505139

**Authors:** Manik Kuchroo, Danielle F. Miyagishima, Holly R. Steach, Abhinav Godavarthi, Yutaka Takeo, Phan Q. Duy, Tanyeri Barak, E. Zeynep Erson-Omay, Scott Youlten, Ketu Mishra-Gorur, Jennifer Moliterno, Declan McGuone, Murat Günel, Smita Krishnaswamy

## Abstract

Biological networks operate within architectural frameworks that influence the state and function of cells through niche-specific factors such as exposure to nutrients and metabolites, soluble signaling molecules, and direct cognate cell-cell communication. Spatial omics technologies incorporate environmental information into the study of biological systems, where the spatial coordinates of cells may directly or indirectly encode these micro-anatomical features. However, they suffer from technical artifacts, such as dropout, that impede biological discovery. Current methods that attempt to correct for this fail to adequately integrate highly informative spatial information when recovering gene expression and modelling cell-cell dynamics *in situ*. To address this oversight, we developed spatial Affinity-graph Recovery of Counts (spARC), a data diffusion-based filtration method that shares information between neighboring cells in tissue and related cells in expression space, to recover gene dynamics and simulate signalling interactions in spatial transcriptomics data. Following validation, we applied spARC to 10 IDH-mutant surgically resected human gliomas across WHO grades II-IV in order to study signaling networks across disease progression. This analysis revealed co-expressed genes that border the interface between tumor and tumor-infiltrated brain, allowing us to characterize global and local structure of glioma. By simulating paracrine signaling *in silico*, we identified an Osteopontin-CD44 interaction enriched in grade IV relative to grade II and grade III astrocytomas, and validated the clinical relevance of this signaling axis using TCGA.

## Introduction

Tissue architecture and cellular spatial dynamics play a significant role in regulating biological systems. However, until recently our ability to measure these dynamics has been limited (1–3). Spatially-resolved transcriptomics is extraordinarily powerful as it provides physiological *in situ* tissue-organizational context to gene expression analysis. While significant, this additional information is accompanied by a high degree of technical noise and artifact, leading to incomplete profiling of cells and difficulties with downstream analysis and interpretation. Most imputation techniques that attempt to recover expression dynamics take into account the gene expression of a cell, but not its spatial location (4–9). A cell’s spatial location, however, provides a great deal of information about its expression properties, as cells occupying the same spatial neighborhood are subject to the same micro-environmental factors and thus often share molecular phenotypes (10). While some recent algorithms have incorporated this information to identify multicellular circuits or enhance resolution (11–13), no technique currently corrects technical artifacts and dropout to identify gene-gene relationships and signaling dynamics free of cell type assumptions. With these insights, we developed spatial Affinity-Based Reconstruction of Counts (spARC), a diffusion geometric framework that integrates *in situ* location and gene expression information to denoise spatial transcriptomic data and identify paracrine receptor-ligand signaling interactions between cells within their spatial contexts. To test the power of this method, we generated the first spatial transcriptomic atlas of human glial tumors and utilized spARC to systematically denoise these data, allowing us to uncover novel pathological cell-cell signaling dynamics.

## Results

### spARC algorithm overview

spARC builds on the connection between manifold geometry, data diffusion and graph signal processing (4, 15, 16) to denoise spatial transcriptomics data. While spatial transcriptomics technologies produce high dimensional data with tens of thousands of genes measured in potentially hundreds of thousands of cells, cellular states are constrained by biological regulatory mechanisms and microenvironmental factors that create interdependencies between genes and spatial neighborhoods (10, 17). Because of these interdependencies, the relevant biological phenomena can be understood in a much lower-dimensional space - or “manifold” - than the dimension of the raw data.

Diffusion maps have been employed as a tool to learn this low-dimensional manifold by simulating a *t*-step random walks along the nodes of the affinity graph (15). To accomplish this, a graph is first computed where nodes represent data points and edges between nodes represent relatedness, estimated by a non-linear kernel function which learns local relationships between data points). Such graphs have been previously computed from single cell data, encoding cells as nodes and cell-cell relationships as edges with varying affinities, and have achieved great success in visualization (15, 18, 19) and clustering (20). In order to learn more global relationships between data points or cells, diffusion maps normalize these affinities into a Markov transition matrix or *diffusion operator* which, when powered to *t*, simulates a *t* step random walks over the nodes of a cellular graph. This effectively captures both long- and short- range relationships within the data. Although initially created for the purpose of manifold visualization (15, 18), diffusion operators can also be applied as a graph filter to denoise (4) and cluster (21) data. Furthermore, recent advances have shown that diffusion operators computed from multimodal datasets (data gathered on the same system by multiple separate sensors) can be merged to produce an integrated diffusion operator that represents the true manifold geometry of the underlying sample (22,23). The technical challenge of multimodal data integration is exemplified by spatial transcriptomics data, in which a single, ground-truth manifold is incompletely captured by two measurement modalities: spatial location in tissue **S** and gene expression profile **X**. spARC uses information from both spatial location **S** and gene expression **X** to denoise a gene of interest **y** (Alg. 1).

#### Algorithm 1 spARC Denoising Algorithm

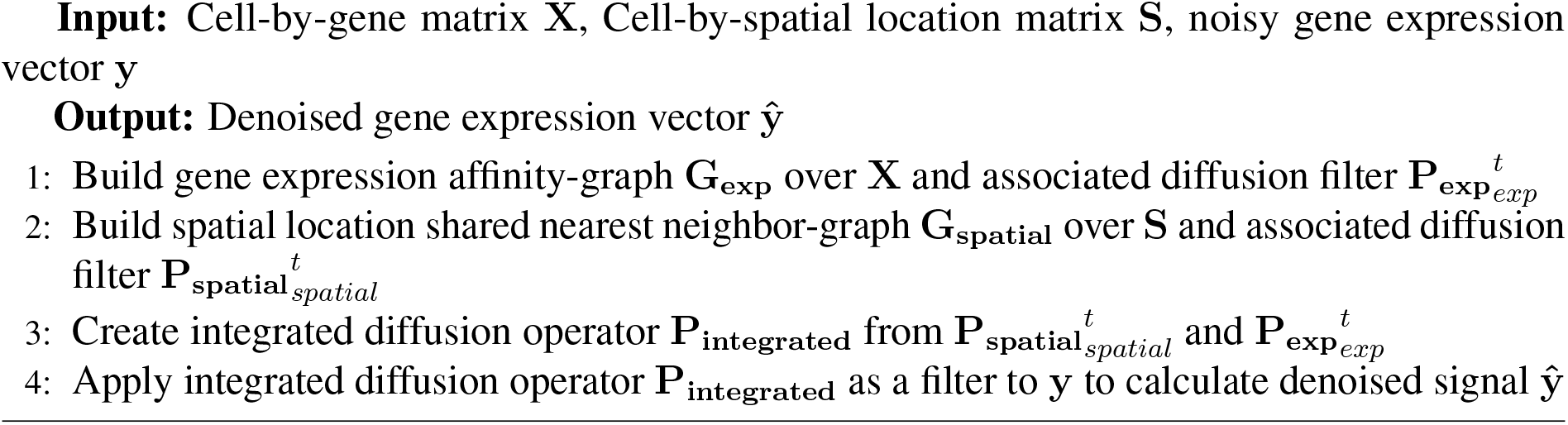

In order to completely model the manifold, we first calculate a nearest neighbor affinity-graph **G_exp_** from gene expression data using **X**, as well as a shared neighbor adjacency-graph **G_spatial_** from each cell’s spatial location using the spatial location matrix **S** (Alg. 1: Steps 1-2) (see Methods). In both graphs, each node represents a cell. In the gene expression graph **G_exp_**, edges represent the similarity in the cells’ measured transcriptomes computed via an *α*-decay kernel as done previously (4, 18, 21). In our spatial graph **G_spatial_**, we would like edges between nodes to not only describe proximity in *in situ* location but also similarity in gene expression; this accounts for divergent cell states within local spatial neighborhoods. For this reason, we construct a shared nearest neighbor graph in which the strength of edges are binary: an edge strength of 1 denotes a relationship that is a near neighbor in *both* the expression values and spatial location, while an edge of 0 indicates a relationship that is *either* not close in spatial location *or* dissimilar in transcriptomic profile. These graphs effectively learn the non-linear dimensions of the single cell data. As described previously, we use these graphs **G_exp_** and **G_spatial_** to compute diffusion operators **P_exp_** and **P_spatial_** respectively through a row-stochastic normalization (see Methods). To learn longer-range relationships in each modality, we raise **P_exp_** and **P_spatial_** to *t_exp_* and *t_spatial_*, respectively, as described previously. Finally, we integrate them into a single diffusion operator **P_integrated_** that represents the spatio-transcriptomic relationships between cells (Alg. 1: Steps 3) and can be used as a diffusion filter to denoise gene expression data in a manner that incorporates both of these information modalities (see Methods).

The diffusion filter has been shown to be effective at learning the cellular manifold and denoising genes based on expression of related cells (4, 22). In contrast to k-Nearest Neighbor smoothing, which averages the expression of similar cells giving them equal weight, a diffusion filter creates a *weighted* average of a gene’s expression based on the strength of edges between cells. In spARC, we use the integrated diffusion operator **P_integrated_**, which contains both spatial and expression information, as a diffusion filter to produce a denoised gene expression vector ŷ (Fig. 1a) (Alg. 1: Steps 4) (see Methods). This process effectively shares expression information between cells with similar transcriptomic profiles and spatial locations via the strength of edges in **G_spatial_** and **G_exp_** respectively. This can theoretically be done by successively applying a diffusion filter directly to the eigenvalues of the Laplacian of each graph; however, to increase computational efficiency we simply multiply the spatial gene expression matrix first by **P_integrated_** as has been done previously (4) (see Methods). In practical terms, this allows a user to analyze a standard Visium 10X sample in under a minute using a freely available Google colaboratory notebook.

**Figure 1:**
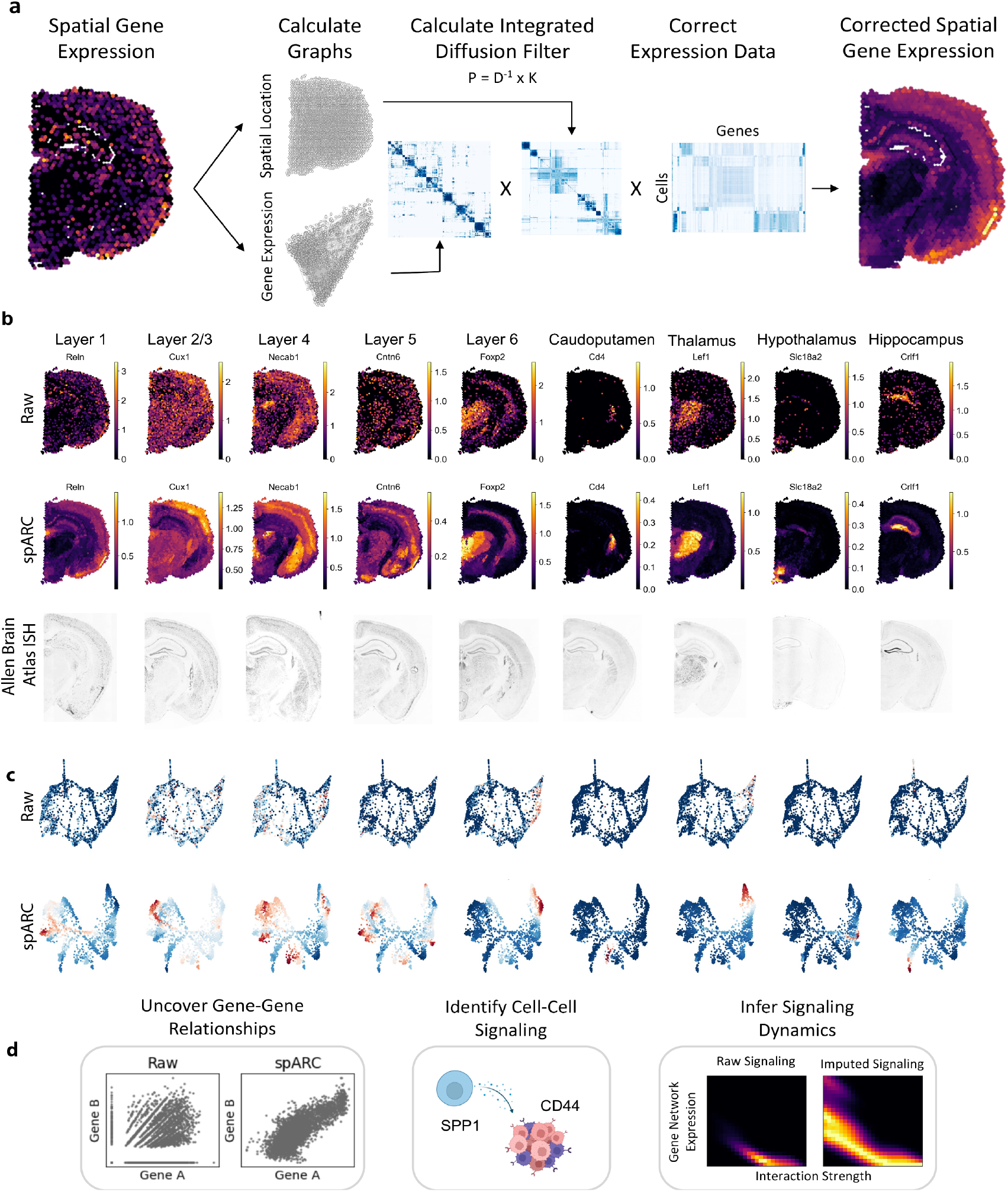
spARC denoises spatial gene expression data to recover gene expression dynamics and signaling interactions. a. spARC computes graphs from both spatial location and gene expression to produce an integrated diffusion filter that can denoise the expression of a gene of interest. b. spARC recovers the expression of key layer-specific genes from the mouse brain measured with spatial transcriptomics 10X Visium platform. These gene expression profiles are validated by ground truth in mRNA situ hybridization (ISH) from the Allen Mouse Brain Atlas (14). c. Visualization of key layer-specific genes in both standard (Raw) and spARC PHATE embeddings. d. spARC resolves gene regulatory patterns based on co-expression patterns, identifies cell-cell signaling interactions and enables inference of signaling dynamics.

We validate that spARC is able to recover ground truth gene expression measurements from spatial transcriptomics data by isolating genes specific to particular layers in the mouse cortex and visualizing raw expression values in tissue measured with 10X Visium platform including *Reln* in layer 1 neurons, *Cux1* in layer2/3 neurons, *Necab1* in layer 4 neurons, *Cntn6* in Layer 5, *Foxp2* in Layer 6, *Lef1* in the Thalamus, and *Crlf1* in the hippocampus (24–30). For many of these genes, we observe minimal layer specificity and little to no correlation between raw transcript measurements and ground truth mRNA (via *in situ* hybridization (ISH) visualization from the Allen Brain Atlas (14)). When spARC is applied, however, gene expression patterns more closely mimic ISH from the Brain Atlas (Fig. 1b). By recovering gene expression patterns, we can begin to study more sophisticated *co-expression* dynamics between cells and simulate the diffusion of ligands in tissue (i.e., paracrine interactions) to infer signaling dynamics between spatially distinct regions (Fig. 1c). We will discuss these applications in the following sections.

### Comparison of spARC to other methods

To benchmark our approach, we compared the performance of spARC to multiple spatial transcriptomic algorithms in recovering ground truth from spatial expression data corrupted *in silico* with dropout and statistical noise (i.e., successful denoising). We opted to carry out this evaluation on data gathered via MERFISH due to its high fidelity and resolution in quantifying transcript counts within their spatial context (31). We simulated increasing amounts of random Gaussian noise and dropout onto expression values (Fig. 2a). We carried out an ablation study comparing spARC to previously-established graph-based denoising techniques (kNN smoothing and MAGIC on the gene expression graph), novel spatial neighborhood denoising techniques (kNN smoothing and MAGIC on the spatial graph), and an integrated approach (kNN smoothing on expression and spatial graph). Across all comparisons and all noise levels, spARC denoising expression values correlated with ground truth expression values far more significantly than with any other approach. After spARC, kNN smoothing that utilized the joint graph (defined over spatial location *and* gene expression) performed best; however, denoising with just spatial location, using either kNN smoothing or MAGIC, performed the worst. This highlights the strength and biological fidelity of integrating both spatial location and gene expression when performing denoising tasks (Fig. 2b).

**Figure 2:**
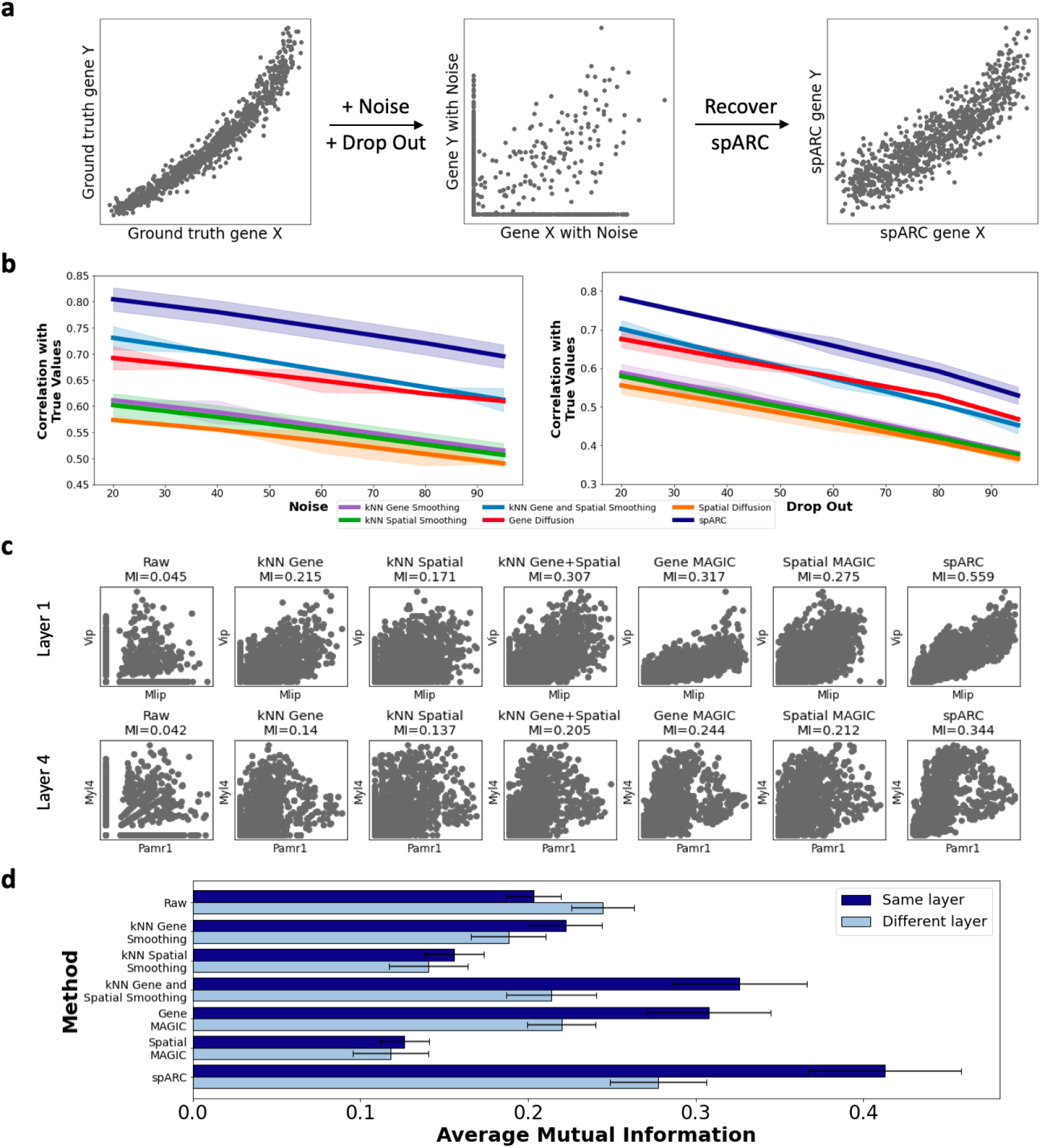
Comparison of spARC with other denoising approaches on spatial expression data. a. Model of simulated noise and drop out in MERFISH spatial transcriptomics data (31). b. Quantitative study comparing denoised results from spARC and other imputation approaches on noisy MERFISH data (31) produced as described in a. Denoised results are compared with ground truth (noiseless) expression values using Pearson correlation as a measure of performance. Experiment is repeated 10 times, shading represents one standard deviation around mean correlation score for each algorithm in each comparison. c. Correlating known co-expressed genes found in layers 1 and 4 of mouse brain after applying different denoising strategies. Mutual information between genes is used as a performance measure, indicating the degree to which gene co-expression dynamics have been recovered. d. Mutual information found between genes known to be co-expressed in the same layer (true association) and genes known to be expressed in different layers (false association) are measured after applying different denoising strategies. This experiment is repeated for 10 genes in each layer, error bars indicate one standard deviation around mean mutual information for each set of comparisons.

We next sought to determine if we could recover interactions between known pairs of co-expressed genes in the Visium mouse cerebral cortex dataset. Using the Allen Brain Atlas "Differential Gene Search” as a reference comparing the layer of interest to pan grey matter, we used *Vip* and *Mlip* expressed in layer 1 and *Pamr1* and *Myl4* in layer 4 (14). Using mutual information, a quantification of non-linear association between two variables, as our performance measure, we found that spARC best recovers gene co-expression dynamics when compared to other denoising approaches (Fig. 2c) (32). To test this exhaustively, we curated a list of 60 genes across 6 layers of the mouse brain that are known to be expressed in a layer-specific manner. By computing mutual information between genes within layers, we were able to test the ability of spARC to recover known layer-specific gene co-expression dynamics better than previous approaches. spARC correctly uncovers known gene programs with greater accuracy (t-test p-value <.05) better than either raw data or other spatial denoising methods, but also out performed MAGIC as well as spatial and gene graph-based kNN denoising methods. While a critical benchmark metric of this work is recovery of co-expressed genes, we do not want to artificially inflate association between genes that are not co-expressed. To ensure that spARC did not significantly produce aberrant gene expression features that do not biologically exist, we computed mutual information between genes expressed in different layers after denoising with each strategy. With this analysis, we find that there is no significant increase in average mutual information found when comparing genes that are not co-expressed when compared to the raw data or other top performing denoising strategies (Fig. 2d) (pairwise t-test p-value>.05). We repeated this analysis using Visium human lymph node data, visualizing genes that help identify the location of key cell types (Extended data fig. S1a). Furthermore, spARC was able to recover known cell type specific co-expression gene expression dynamics better than other approaches while not falsely inflating background noise (Extended data fig. S1b,c).

To further confirm that spARC recovers both shared and divergent transcriptional states in the local tissue microenvironment, we turned to spatial transcriptomic profiling of human lymph node. Within secondary lymphoid tissues immune cells are organized into discrete compartments. Antibody-producing B cells reside in follicles, typically near the outer capsule where antigens enter the tissue (33). During an immunological insult, B cell activation drives a sequence of migratory events that ultimately leads to formation of micro-structures within follicles called germinal centers (Extended Data Fig. S1d). The architecture of the germinal center (GC) begets a classical Darwinian evolutionary process, in which antibody-producing B cells undergo spatially separate phases of diversification, followed by selection on the basis of antibody-antigen binding affinity, ultimately improving the ability of the host immune system to target a given pathogen with high-affinity antibodies following immunization or infection. GCs are organized into two functionally distinct regions; in the dark zone (DZ), B cells undergo programmed somatic mutation of genomic regions that encode antibodies. They then migrate to the light zone (LZ), where they compete for pro-growth signals delivered via cognate interaction with CD4+ T follicular helper (TFH) cells. The incidence, duration, and strength of these interactions is dictated by presentation of antibody-captured antigen to TFH cells, therefore it is directly proportional to antigen affinity and allows for selection of high-affinity antibody producing cells. B cells undergo many rounds of iterative mutation and selection, generating high-affinity antibody producing cells that can seed long-lived memory subsets and provide lasting immunological protection (33, 34).

We first investigated whether spARC was able to improve visualization of these structures and their sub-compartments. LZ and DZ gene expression signatures have been established prior to spatial single cell technologies through experiments where cells in either the light or dark zone were spatially labeled *in situ* by two-photon activation of a transgenic photoactivatable-GFP reporter protein, then sorted and RNA-sequenced in bulk *ex vivo* (35). Using these gene signatures (LZ: *CD19*, *CCND2*, *IL4I1*, *CD40*, *CD83*, *MYC*, *CD86*, *AICDA*), (DZ: *CD19*, *CCNB2*, *HMMR*, *POLH*, *LMO4*, *CXCR4*, *CCNA2*), we show that spARC recovers the separation of TFH-adjacent LZ and TFH-distal DZ compartments (Extended Data Fig. S1e) and expected frequencies of LZ and DZ B cells (Extended Data Fig. S1f). Within the LZ, cells that are undergoing TFH-mediated positive or negative selection occupy the same anatomical niche, yet have polarized transcriptional programs. This provides a model opportunity to investigate the extent to which spARC recovers distinct transcriptional programs in nearby cells. Gene expression of B cells actively receiving or not receiving T cell help has been experimentally determined (36) and was used to visualize transcriptional separation of LZ B cells (Extended Data Fig. S1g). spARC recovers two distinct populations within the LZ that express either genes induced by TFH (*CCND2*, *BATF*, *JUNB*, *IRF4*, *MYC*, *CD69*), or genes associated with negative selection or steady-state (*BTG1*, *BTG2*, *POLH*, *NR4A1*, *CD72*, *BACH2*, *FOXO1*, *KLF2*, *IRF8*), showing that spARC is able to differentiate divergent transcriptional states in local cell populations.

Finally, we show that our flexible framework is extendable to any number of data modalities. Using DBIT-seq data, which measures protein and gene expression *in situ* (37), we find that we can recover associations between key marker genes and their translated protein counterparts significantly better than other approaches (Extended Data Fig. S2)a,b. We applied spARC to a range of spatial technologies, including slide-seq (38, 39), seqFISH (40), MERFISH (31), MIBI-TOF (41) and 4i (42), and find that spARC is generalizable across platforms (Extended Data Fig. S2 c-g).

### spARC simulates paracrine signaling interactions

#### Algorithm 2 spARC Interaction Algorithm

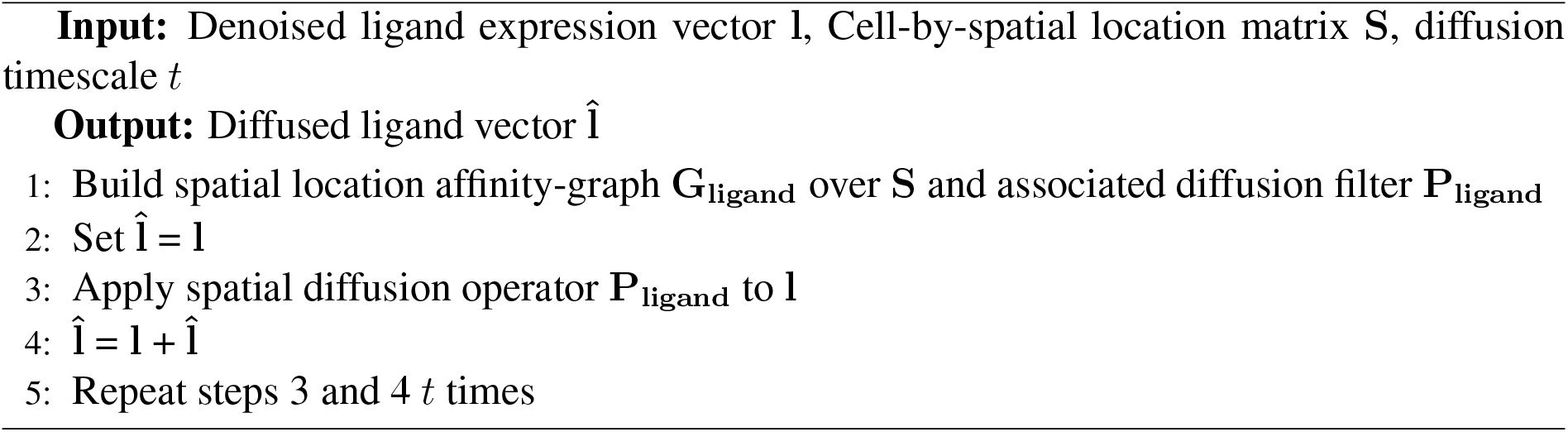

A key advancement of high-dimensional spatial transcriptomics is the integration of spatial context with potential gene-gene signaling networks, or *gene programs*. Previous approaches to discovering gene programs *in silico* define clusters based on transcriptome data alone and compute interaction “scores” for ligand-receptor pairs using expression levels per cluster; these do not incorporate spatial information (11, 43–45). While informative, this not only fails to account for proximity of cells, which physically bounds the set of possible receptor-ligand interactions as they have a limited diffusion radii, but also relies on user-defined clusters. An ideal approach would simulate signaling interactions between individual cells, free of any cluster assumptions.

Conveniently, the diffusion operator constructed using a Gaussian kernel also corresponds to an exponentially decaying concentration gradient for a molecule originating from a single source (this is simply the solution to the 2-dimensional diffusion equation, except we now have a discretized space). Thus, we can model the physical diffusion of a ligand through space using the same set of transition probabilities as in our diffusion operator. Since we are mainly interested in gene programs, or relationships *between* genes, we do not need the actual chemical concentrations, and can simply use RNA expression level as proxies for the *relative* presence of multiple different ligands.

We then simulate the diffusion of ligands in tissue by applying the spatial diffusion operator (Extended Data Fig. S3a). In this setting, we compute a new spatial nearest-neighbor affinity graph **G_ligand_** (Alg. 2: Steps 1), its associated spatial diffusion operator **P_ligand_** (Alg. 2: Steps 2), and serially apply this operator to the denoised ligands **l** to simulate its diffusion in physical 2-dimensional space (Alg. 2: Steps 4-5). This process simulates the concentration gradient of a ligand of interest **l** across *t* different diffusion timescales (46)(Extended Data Fig. S3b). By combining these diffused values with the denoised expression of cell’s cognate receptor, we can compute paracrine signaling scores. Unlike previous methods (43, 47) which map interaction strengths with clusters of cells, spARC is able to recover and simulate single-cell interaction strengths, allowing for downstream correlation analyses to identify both known and novel gene programs.

### Comparison of spARC interaction analysis

To test spARC’s ability to recover known cell-cell interactions *in situ*, we first simulated the diffusion of two ligands, IL1B and IL6 in our 10X Visium lymph node dataset. Using known ligand-receptor interaction databases, we found the expression of their cognate receptor and computed ligand-receptor interaction scores per cell (43, 47) (see Methods). As ligand-receptor interactions often produce downstream gene expression effects on the cognate cell, we can correlate these single-cell interaction scores with known IL1B and IL6 signalling response genes (48). From these comparison, we can clearly see that IL1B and IL6 signaling response signatures overlap a significant amount with our spARC computed interaction scores (Extended Data Fig. S3b and c). In order to compare spARC’s performance to other established algorithms (43, 49), we compared the signaling interaction score for IL6 and IL1B using each algorithm and associated these scores with IL6 and IL1B signaling response programs using mutual information as a performance measure. Across both comparisons, spARC had significantly higher association between computed interactions scores and downstream signaling signature (Extended Data Fig. S3d and e). For a more systematic comparison spARC to these methods, we calculated 634 distinct ligand-receptor interactions in each cell in the lymph node dataset and associated these interaction strengths with established ligand-receptor response gene signatures (48). We saw that across all comparisons, spARC significantly outperformed both CellPhoneDB and SpaOTsc at matching interaction strengths to downstream gene expression programs associated with these interactions (Extended data Fig. S3).

Finally, there are other uses of a diffusion operator including visualization and clustering. While we do not expand on this topic here, we do show that visualizations of this spatial transcriptomic integrated diffusion operator can be clustered and visualized like any single modality diffusion operator and in fact, can reveal spatial organization better than established dimensionality reduction approaches (Extended Data Fig. 1) (18).

### spARC identifies spatial niches and signaling interactions implicated in glioma progression

We then used spARC to study networks and signaling relationships in IDH-mutant (IDH-MUT) glioma tumors across grades. The prognosis of Glioblastoma (GB) is grim, with the median survival only marginally improving from 10.1 months to 14.6 months in over 100 years, despite numerous advances in therapeutic, surgical, and radiological techniques (50). Using WHO 2016 criteria, primary GB can be classified into *IDH* wildtype (IDH-WT) or *IDH* mutant (IDH-MUT) types, with secondary GB almost exclusively arising from lower grade IDH-MUT gliomas. (51). These IDH-MUT tumors can be further classified into IDH-MUT with 1p/19q co-deletion indicating its transformation from oligodendroglioma, or IDH-MUT astrocytomas which commonly have *ATRX* and *TP53* mutations (52). While IDH-MUT gliomas have better survival than IDH-WT, the population that is most affected is significantly younger (53).

We applied spARC to 25,517 barcoded array spots from 10 surgically resected human gliomas: WHO Grade II (n=3) and WHO Grade III (n=3) and (WHO Grade IV (n=4). In order to study signaling co-expression networks across disease progression, we chose to only study IDH-MUT astrocytomas, 9 of which had *TP53* and *ATRX* co-mutations (Extended Table. S1). Next, we performed denoising of each sample independently with spARC and visualized gene programs across the samples including dense tumor, infiltrating/high tumor burden, leading edge, hemoglobin, and vasculature, described in the next section (Extended Data Figs S4).

One sample, GB-1182, captured a region of histological heterogeneity with focal solid dense tumor surrounded by hypercellular brain parenchyma diffusely infiltrated by tumor, proliferated blood vessels and regions of extravasated blood. Using diffusion condensation (54), a graph clustering method we previously developed, we identified five clusters which significantly overlapped with histopathological annotations (Fig. 3a). The annotated region of dense tumor had high expression of known tumor associated genes *CD44*, *VIM*, *LHDA* (Fig. 3b). Pathway enrichment analysis of the top 100 genes for this cluster revealed an enrichment in Plasmin Effects in Inflammation (p-adj = 1.851e-11) and Glycolysis Activation in Cancer (Warburg Effect) (p-adj = 6.229e-11) (Extended Data Fig. S6a). Additional analysis of this pathway was obtained by performing our previously published method conditional-Density Resampled Estimate of Mutual Information (DREMI) on the matrix product of these genes to identify specific areas of co-expression (Extended Data Fig. S6a) (55). This approach identified additional Plasmin-related genes with spatially restricted expression including immediate early genes (*FOSB*, *IER2*), and genes downstream to plasmin activation (*CCN1*) and invasiveness in glioma (*LCTL*) (Extended Data Fig. S6b) (56–58). Separately, spatially localized expression of *ERO1A*, *VDAC2*, *ADM*, and *VEGFA* were among the glycolysis and angiogenesis related genes identified when applying DREMI to the Glycolysis Activation in Cancer (Warburg Effect) gene module (Extended Data Fig. S6c) (59, 60). Both of these processes represent fundamental tumor promoting programs of hypoxic metabolism and promotion of tumor angiogenesis. This approach highlights the potential for unbiased marker gene discovery using spatial transcriptomics.

**Figure 3:**
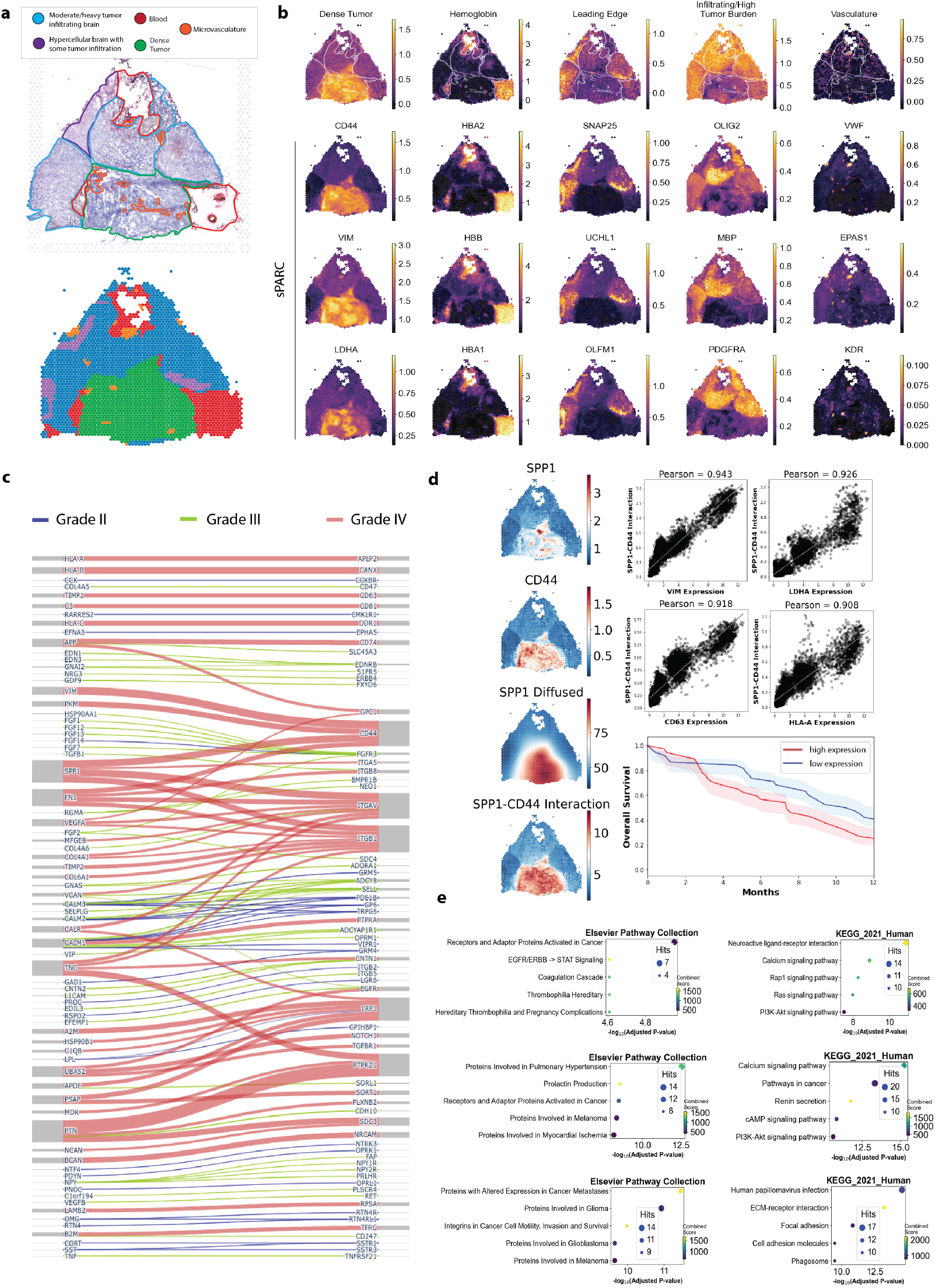
spARC identifies tumor subtype specific regulatory networks from spatial glioma transcriptomics data. a. Annotated pathology of GB-1182 demonstrating 5 major characteristics in the histology. Below, diffusion condensation cluster map of sample. b. Tumor niches described as gene modules including Dense Tumor, Hemoglobin, Leading Edge, Infiltrating/High Tumor Burden, and Vasculature. Representative composite image is on the top row with annotation outline overlaid. Representative top genes shown below with sPARC imputation. c. Network graph of receptor-ligand interactions of top 50 genes that are enriched in each grade relative to all other samples. Blue is Grade II, Green is Grade III, and Pink is Grade IV. d. Expression of each gene (SPP1 and CD44), simulated diffusion of SPP1 and overlapping region of interaction of SPP1-CD44. Selection of top pearson correlated genes to the receptor-ligand interaction. Gene co-expression of SPP1 and CD44 is used to predict outcome in TCGA data. e. Elsevier Pathway Collection and KEGG 2021 Human enrichment is shown for each analysis. Top) Grade II, Middle) Grade III, Bottom) Grade IV.

In agreement with the histopathological annotation, hemoglobin genes *HBA2*, *HBB*, *HBA1* were highly expressed in regions belonging to blood and large blood vessels (Fig. 3b). There were two distinct regions in the infiltrated brain. The first region was enriched with mitochondrial genes and genes known to be important to GB biology including *OLIG2*, *BCAN* and *PDGFRA* (61). Enrichment analysis of this region revealed two distinct clusters with spatially restricted gene expression: Glioblastoma, Proneural subtype (p-adj = 5.7e-5, Extended Data Fig. S6d) with top DREMI-ranked genes including *OLIG2*, *COL20A1*, *MAP2*, and *SOX2* (Extended Data Fig. S6e). The other scluster was enriched for Astrocyte Dysfunction and GABA Signaling Deficiency (p-adj = 1.3e-3, Extended Data Fig. S6d) with top DREMI-ranked genes including *AQP4*, *SLC1A3*, *SLC39A12*, and *SERPINA3*. Notably, genes in this gene module overlapped with the histopathological annotation of *hypercellular brain with tumor infiltration* (Fig. 3a & b). The second region in the infiltrated brain, *Leading Edge cluster*, appeared at the border between the tumor and brain tissue and showed a high degree of overlap with previously defined genes belonging to the Leading Edge according to the Ivy GB database (Extended Data Fig. S6g & i) (62). The top gene in this region was *SNAP25*, which was a gene previously shown to be specific to leading edge by Ivy GB database and subsequently identified as inversely related to glioma progression (63). Upon enrichment analysis, *SNAP25* was found in several of the enriched terms including Exocytosis:Vesicle Fusion (Extended Data Fig. S6g). Top DREMI-ranked genes in this region included additional genes related to neurotransmitter release (*MCTP1*) and glioma progression (*LY6H*) (64, 65). The final region observed was determined to be vasculature, which demonstrated well-known vascular genes such as endothelial markers *KDR*, *EPAS1*, *VWF*, *PECAM1*, as well as smooth muscle marker *CDH5*. All these regions were seen to various degrees between our samples of other GB and WHO grade II and grade III tumors (Extended Data Fig. S5).

To understand interactions between regions of tumor in 2D space, we simulated ligand diffusion (see Methods). By identifying diffused ligands from a curated database and performing differential expression, we identified gene interactions between previously established receptor-ligand pairs across all grades (Fig. 3c) (43). First, we wanted to determine signaling pairs that were enriched in GB samples with higher grades relative to lower grades. Using the top 50 interaction pairs, we identified that signaling involving *CD44*, *SPP1*, *FN1*, *ITGB1*, *ITGAV*, *LRP1*, and *PTPRZ1* occurred in more than 3 receptor-ligand interactions. Among these interactions was SPP1-CD44, which has been previously characterized to play an important role in high-grade glioma and GB (66, 67). We also identified receptor-ligand interactions that were enriched in WHO Grade III and WHO II (Extended Data Fig.3c & e)

In order to identify the genes most regulated by this interaction, we computed the top 10 genes whose expression is correlated with the SPP1-CD44 signaling interaction. Among the genes potentially regulated by SPP1 signaling were *VIM*, *ANXA2*, and *CD99*, all previously implicated in regulating aggressive behavior in high grade gliomas and GBs (67–69). Other genes include *A2M*, *S100A10*, *C1R*, *RASD1*, and *CRYAB*. Mapping these top 10 genes onto survival data produced by TCGA, we identified that patients with tumors that possess a higher expression of these genes have significantly greater mortality than patients that express lower levels of this signature, further confirming the pathogenic nature of SPP1-C44 signaling axis (Extended Data Fig. 3d). In GBs as well as other tumors, SPP1-CD44 signaling axis has been shown to play a role in stem cell maintenance and immune infiltration. By examining the top 100 genes correlated with this receptor-ligand interaction we found genes associated with hypoxia regulation *LDHA*, and immune markers *CD63*, *HLA-A*,*HLA-B*, *HLA-DRB1*, *TNFRSF1A*, and *LGALS1* (Fig. 3d). Enrichment analysis of these 100 genes using KEGG Human 2021 revealed that the top term was antigen processing and presentation (p-adj = 9.9e-10). While the SPP1-CD44 signaling axis has been recognized in gliomas and GBs, prognostic value has been primarily attributed to CD44 expression (70). The *in situ* identification and resolution of signaling partners within spatial compartments gives further evidence of the importance of this axis in GBs and demonstrates spARCs power to give broader insight into signaling cross-talk.

## Discussion

Spatial transcriptomics offers powerful tools to study *in situ* gene expression and model gene interactions. However, like other whole-transcriptome assays before it, its utility is hindered by high degree of dropout and noise. Current imputation and denoising techniques that attempt to recover expression dynamics only take into account a cell’s gene expression (4–9), overlooking the fact that spatial location provides a great deal of information about state and function. Proximal cells are exposed to the same microenvironmental factors such as nutrients, soluble signaling molecules with a limited diffusion radius, and contact-dependent ligand-receptor interactions with other cells. By partially encoding these extrinsic features, the spatial location of a cell is informative in ways not captured by transcript or protein measurements alone.

Recently, problems in exploratory analysis of spatial omics data have been approached successfully with methods that incorporate spatial information. For example, DIALOGUE (11) identifies cellular interaction circuits by initially subsetting cell types based on shared gene expression and shared spatial location. SpiceMIX (12) identifies cell types and spatially variable genes by employing a graphical representation of the spatial arrangement of cells. Downstream analysis, however, is bottlenecked by the quality of upstream data processing, an issue spARC aims to address. These problems could be well-served by incorporating spatial information in a *supervised* way, as demonstrated by BaySpace (13), which enhances resolution of spot-based spatial transcriptomics by clustering spot-measurements with an imposed spatial proximity *prior*. However, to our knowledge, the major issue of technical noise and dropout has yet to be directly addressed with combined spatial and gene expression information in an entirely unsupervised manner. In this manuscript, we develop a tool that does just this. We leverage combined spatial and gene expression features with a novel machine learning algorithm that uses manifold learning, graph filters, data diffusion and diffusion filtration to enable discovery of gene-gene relationships, spatial co-localization of genes, and receptor-ligand interactions applicable across a variety of spatial transcriptomic methods.

We first applied spARC to denoise mouse brain transcriptomics and were able to recover ground-truth in situ hybridization mRNA expression that enabled the specific visualization and analysis at the level of anatomical structures such as the restricted expression of layer specific genes in mouse brain. Furthermore, we uncovered receptor-ligand interactions from 10 surgically resected human gliomas (WHO Grade II (n=3), WHO Grade III (n=3), and (WHO Grade IV (n=4). Through this analysis we found specific genes modules such as the Leading Edge, which were consistently identified in tumors adjacent to infiltrated brain. Not all of the samples showed the same degree of restricted spatial localization and the difference was most obvious between samples with tumors adjacent to other tissue versus those where the sample was uniformly tumor. The observation of a distinct spatial transcriptome at the tumor-tissue interface has been recently noted in an *in vivo* model and spatial analysis of melanoma (71). This may be explained by considering that locality dictates factors such as nutrient availability, access to soluble signaling molecules with a limited diffusion radius, and direct contact with proximal tumor cells. Unbiased exploration of the differences between the tissue-tumor interface compared to the tumor core may enable a better understanding of the ways that GBs are able to resist multiple treatment modalities and identify expression signatures amenable to targeted treatment.

Spatial sequencing enables the *in situ* assessment of pathogenic and developmental states that have until now remained unobserved. The future application of these methods with previously established computational histology, time-series experiments, and perturbations could greatly enhance our understanding of two-dimensional spatial patterning and tissue morphogenesis in physiologic and pathologic conditions (72).

## Acknowledgments

We would like to thank the donors and their families for their contribution to this work. Without their sacrifice, our study would not have been possible. We would also like to thank Guilin Wang PhD, Christopher Castaldi MS, Kaya Bilguvar MD, and the Yale Center for Genome Analysis for their support.

This work was supported by NIAID training grant 1F30-AI157270 (to M.K.) and by NIAID 5U19-AI089992-08 (to S.K.), NIGMS 1RO1-1355929 (to S.K.), Gregory M. Kiez and Mehmet Kutman Foundation and Yale School of Medicine funds (to M.G), NIH/NCI no.F31CA254426 (D.F.M), NIH-Medical Scientist Training Program no.T32GM007205 (M.K. and D.F.M.), and Connecticut Brain Tumor Alliance (J.M. and M.G.)

## Methods

### Genome Sequencing, Tissue Preparation, Cryosectioning and Imaging for Visium 10x Spatial Gene Expression

Patients at Yale New Haven Hospital undergoing neurological surgery were consented for sample collection. All samples are from patients diagnosed and operated on between 2016-2018 and were classified according to 2016 WHO guidelines at time of diagnosis. DNA was extracted from tumor and blood samples and sent to Yale Center for Genome Analysis for whole-exome sequencing. Data was analyzed using previously published methods from the Günel lab (73) and quality control metrics can be found in Supplementary Table 1 S1. Portions of tumors were immediately flash frozen as blocks in in OCT and stored at −80°C until spatial preparation. To prepare for use with 10x Visium Spatial Sequencing, OCT blocks were sectioned into 10μm slices and immediately placed on Visium array slides (Visium Spatial Gene Expression slides, 10× Genomics). Hematoxylin and Eosin (H&E) staining, RNA extraction, and cDNA strand synthesis was followed as directed in the Visium protocol (https://support.10xgenomics.com/spatial-gene-expression). Images were captured using the Keyence BZ-X700 microscope system using 40x magnification. Sequencing was performed at Yale Center for Genome Analysis and Visium samples were sequenced to 75 million pairs/sample. Transcriptome alignment and counts were generated using 10x Space Ranger version 1.3.1.

### Validation and Pathological Review

For imputation expression validation, several methods were employed. First, we used ISH generated by Allen Mouse Brain Atlas (14) to identify genes that are specific to anatomical brain locations in mouse. Coronal images were then imported into Adobe Photoshop, cropped to only include half of the cortex, and individual adjustments to whole picture brightness, contrast, and saturation were applied to achieve a representative image of the gene expression. All original images can be viewed interactively at mouse.brain-map.org using the gene name indicated in the figure. For lymph node protein expression, frozen splenocytes from C57BL/6N mice (Charles River) were thawed at 37°C and washed in 1XPBS with 1% Bovine Calf Serum (Sigma F8192) and 2mM EDTA. Single cell suspensions were labeled for 20 minutes on ice or 4°C for 20 minutes with 10% Rat serum (STEMCELL Technologies), Fixable Viability Dye at 1:400 (ebioscience 65-0866-18) and surface antibodies CD19 Brilliant Violet 711 (BD Biosciences 63157), CD38 PerCP/Cy5.5 (Biolegend 102721), GL7 FITC (Biolegend 144603), CD86 Pacific Blue (Biolegend 105021), CXCR4 APC (Biolegend 146508) at a final concentration of 1:200. Multi-colour cytometry was performed on the LSR II flow cytometer (BD biosciences) and analyzed with FlowJo v10.6.2. For glioma expression validation, samples with leftover tissue were used to validate the protein expression of key genes OLIG2 (Millipore Sigma AB9610), GFAP (Invitrogen PA110019), and KI-67 (Abcam ab15580). Briefly, standard immunohistochemistry protocols were employed for fresh frozen tissue which included an H202 block, 2% block BSDSGS (Jackson ImmunoResearch) with 0.3% triton followed by overnight incubation with an antibody concentration of 1:250 and secondary antibody (Jackson ImmunoResearch) in 2% BSDSGS at 1:250 concentration. Substrate reaction was performed using Vectastain ABC kit (Vector Laboratories PK-4000) and ImmPACT® DAB EqV Peroxidase (HRP) Substrate (SK-4103). H&E stained images used for Visium Spatial Gene Expression were reviewed by a board-certified neuropathologist (D.M.) and annotated for cellularity, mitoses, necrosis, microvascular proliferation. Annotations were performed blinded to the results of clinical/pathological diagnosis and to our computational analysis.

### Computational Methods

In the following sections we provide a thorough background on manifold learning, graph filters, data diffusion and diffusion filtration as well as a description of each aspect of the spARC algorithm, including diffusion filtration-based denoising, shared nearest neighbor graph construction and signaling simulation. Finally, we explain specifics around how signaling and denoising comparisons between algorithms were run.

### Background on manifold learning and graph theory

While single cells are measured in a high number of dimensions, the intrinsic dimensionality, or directions of variation, within a dataset are limited. This is because each gene measured does not behave independently of all other genes. Due to complex and non-linear regulatory mechanisms, these genes are expressed in synergistic or antagonistic manners. These co-regulated modules form fewer axes of variation and can be used to modal a majority of the variability found in the input dataset. A common way of learning the single cell data manifold is by computing a nearest neighbor affinity graph *K* using a Gaussian kernel function:

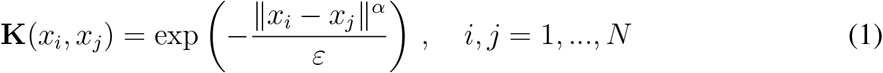

Where **K** is an square *N* × *N* matrix whose (*i, j*) entry, denoted by **K**(*x_i_, x_j_*), emphasizes the similarity between cells *x_i_* and *x_j_*, based on kernel bandwidth parameter *ε*. Recently, alpha-decay kernels have been proposed and used in single cell tool development (4, 18, 21). While a Gaussian kernel function sets *α* to 2, in the alpha-decay kernel, *α* is set to a large positive number, making affinities rapidly diminish between more distant cells. While previously, *x_i_* and *x_j_* have denoted cells gene expression profiles, an affinity matrix can be created using any set of features measured in every cell, including chromatin accessibility, protein expression and spatial location. In each of these setting, high affinities denote cells that are similar to one another and low affinities denote cells that are dissimilar.

### Background on data diffusion and integrating multimodal data

In (74), diffusion maps were proposed as a robust way to capture intrinsic manifold geometry in dataset by eigendecomposing a Markovian random walk matrix that simulates cell state transitions in single cell data. A diffusion operator simulates the diffusion of a signal along the nodes of a cellular graph through *t*-step random walks. This process effectively reveals non-linearities present within single cell data and has been shown to effectively model the single cell manifold (4, 18, 21)

A diffusion operator is calculated by:

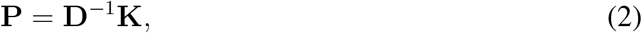

where **D** is diagonal degree matrix and K is the affinity matrix both previously described. The matrix **P**, or diffusion operator, defines the single-step transition probabilities of a Markovian random walk. Intuitively, the eigenvectors and eigenvalues of the diffusion operator **P** are equivalent to those of the normalized graph Laplacian *L* = *I* − *P* = *D*^−1^*L_u_* = *D*^−1^(*D* − *K*), *K* is the kernel affinity, *L_u_* is the unnormalized graph Laplacian, where *I* is the identity matrix, *D* is the degree matrix.

Recently, (23) proposed alternating diffusion as a means to extend the data diffusion approach to combine diffusion operators created from multimodal data. Intuitively, this generalizes the *t*-step random walk to taking alternating steps in each data modality. Practically this is done by taking a product of each diffusion operator **P_a_** and **P_b_**.

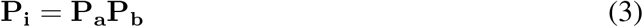

Finally, the resultant alternating diffusion operator **P_i_** is powered to *t* stimulate alternating steps across modalities. (22) extended this framework to include different data denoising tasks before data integration to weight the information in from each data modality separately as they may include differing amounts of information. Experiments revealed that this integration procedure improved data visualization and denoising tasks.

### Background on Graph Signal Processing

Graph Signal Processing (GSP) extends the field of signal processing from the spatiotemporal domain to to data that can be modelled on a graph, such as single cell transcriptomics data. GSP extends this framework to now ‘filter’ signals, or the expression values of a particular gene on the nodes of a graph, to produce denoised signals. In the past this framework has been used successfully in single cell data to denoise individual genes and compute enrichment likelihoods between perturbation conditions (4, 16). All of these graph filtration strategies fundamentally rely on the similarities between classical Fourier transform and graph Fourier transform (GFT) which is derived through eigendecomposition of the graph Laplacian.

The graph Laplacian can be defined as:

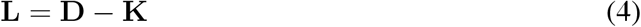

Where L is a square *N* × *N* graph Laplacian matrix and **D** is diagonal matrix: **D**(*x_i_*, *x_i_*) = ∑_*j*_ **K**(*x_i_, x_j_*). The eigenvectors Φ of the graph Laplacian **L** have been shown to be equivalent to graph frequency harmonics (75). Thus, signal loadings on to these eigenvectors creates a *graph Fourier transform* defined as Φ*^T^ f* for a graph signal *f* which can be filtered similar to signals in classical signal processing to produced a denoised signal.

A graph filter itself is a function that modulates the loading coefficients of signal *f* on to the eigenvectors of the graph Laplacian. Thus, a *graph filter h* can be defined as

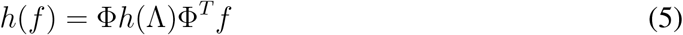

Where eigenvectors and diagonalized eigenvalues of the graph Laplacian **L** are Φ and Λ respectively and h() represents a filter function. Here, the properties of the filter function *h* rescale the eigenvalues of the graph Laplacian to modulate frequency components of *f* to produce denoised signal h(f). Many types of filter functions have been established previously including high pass and low pass filters, however two filters that have gained traction in computational biology field for studying single cell data are the diffusion filter and the related heat filter. Based on the field of data diffusion, we expand on this particular filter in the following section.

### Denoising data with data-diffusion

In (15), powers **P**^*t*^ of the diffusion operator, for *t* > 0, not only simulate *t* step random walks over the data, defining longer range diffusion, but also can act as a filter function, modulating the eigenvalues Λ of diffusion operator **P**. This diffusion filter acts as a soft low-pass graph filters 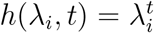, which rapidly diminishes the value of high freqeuncy eigenvectors while only maintaining the low frequency eigenvectors. While a true diffusion graph filter requires eigendecomposition of the **P**^*t*^ matrix, this is computational expensive. Since **P**^*t*^ = ΦΛ^*t*^Φ^*T*^, we can avoid eigendecomposition and simply apply the powered diffusion operator **P**^*t*^ directly to the noisy signal as a *diffusion filter* directly:

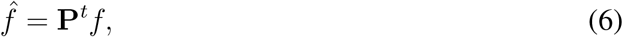

### Denoising spatial transcriptomics data with spARC diffusion filters

spARC integrates these advances in GSP and multimodal data diffusion to denoise spatial transcriptomics data with an integrated diffusion filter. First spARC calculates graphs for all data modalities. For expression modalities, such as gene expression and protein expression, these graphs are alpha-decay affinity graphs 1. The spatial graph however, is constructed using a shared nearest neighbor approach to only maintain connections between cells that are both transcriptomically similar and in the same spatial neighborhood. This graph construction is described below.

After computing each of these graphs, we compute diffusion operators **P_spatial_** and **P_exp_** through row normalization of their respective affinity matrices as in 2. We then power these diffusion operators **P_spatial_** and **P_exp_** to *t_spatial_* and *t_exp_* respectively and integrate them into a single diffusion operator **P_integrated_** as described in (22, 23) and equation 3. Finally, we apply this integrated operator as a filter to denoise the expression of a gene of interest:

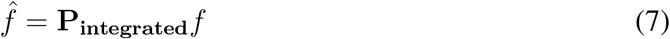

Using default settings, spARC calculates the **G_spatial_** and **G_exp_** the bandwidth *ϵ* of the adaptive alpha-decay kernel is set to the distance to the 15th k-Nearest Neighbors for each point, *t_spatial_* is set to 2 and *t_exp_* is set to 3. These parameters can all be set by users.

### Shared nearest neighbor spatial graph calculation

The need for the shared nearest-neighbor graph arises from a key observation. In the transcriptome graph, for a given number of neighbor of nearest neighbors, these neighbors *also* tend to be spatially co-located. However, on the *spatial graph*, for any fixed number of nearest neighbors, these neighbors are often transcriptomically dissimilar. That is, neighboring cells are often of differing cell types - this is consistent with prior observations of spatial heterogeneity of many cell types. This can be considered a form of “noise” intrinsic to the graph construction which is not conducive to recovering our signal of interest, which is gene expression.

We can see the consequences of this noise this empirically: carrying out MAGIC (a graph denoising procedure) on a naively constructed spatial graph, with a fixed number of nearest neighbors, performs poorly.

To get around this, we make the stringest requirement that spatial graph nodes be connected if and only if they are *also* nearest neighbors in the transcriptomic graph. We also make this *binary*. Formally, for nearest-neighbor diffusion operators defined on the transcriptome domain, **G_exp_** and spatial domain, **G_spatial_**, the spatial graph shared nearest-neighbors graph is computed as:

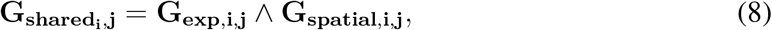

where ∧ denotes the binary intersection operator. In plain English, edges are binary [0,1] depending on if two cells occur in the same spatial neighborhood AND if they are similar transcriptomically. In order to create this graph, we calculate the nearest k neighbors for every cell. If a cell is within these k nearest neighbors in *both* spatial and expression modalities, the edge weight between the two cells is set to 1; if not, it is set to 0. This allows us to create a graph that encodes spatial edges between *similar* cell types.

### Modeling ligand diffusion and paracrine signaling interactions

Modelling realistic paracrine signaling interactions is a key advance made by spatial transcriptomics data. spARC is able to leverage both transcriptomic data and distance information between cells to simulate concentration gradients of ligands as they diffuse in tissue. By modeling the concentration of diffused ligands in tissue and by taking into account receptors of these ligands, we can model signaling interactions and correlate downstream effects.

In order to simulate the diffusion of ligands in tissue first we create a new spatial graph **G_ligand_** as in 1. Importantly, this is an alpha-decay kernel graph and not a shared nearest neighbor graph as the expression of local cells is not meaningful to simulate ligand diffusion. Using this graph we can simulate the diffusion of ligand in tissue by heavily filtering the denoised spARC ligand’s expression:

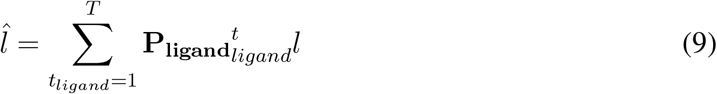

where l is the denoised expression of a ligand, **P_ligand_** is the diffusion operator calculated from **G_ligand_** via 2 and *t_ligand_* is a measure of ligand diffusion distance. At default, spARC set’s T to 10.

Once we have computed this concentration gradient for a ligand, we can compute interaction scores between it’s cognate receptor by multiplying the concentration of the ligand at a location 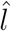 with the expression of the receptor at that location. From here we can compute downstream consequences of signaling using other association strategies such as linear regression, mutual information and DREMI (76). To obtain specific expression of gene-sets and identify co-localized genes, gene matrices were multiplied together, prior to use with DREMI. We have curated a custom receptor-ligand interaction databased by combining the 834 curated ligand-receptor combinations and multi-unit protein complexes found in CellPhoneDB (43) with the 2557 ligand-receptor interactions found in the celltalker database (47).

### Visualizing spatial transcriptomics data with spARC diffusion operator

As described previously, a diffusion operator can be used to visualize (15, 18) complex data. Recently, the PHATE algorithm has built on the data diffusion framework to visualize complex single cell data in 2 or 3 dimensions. Here, we build a PHATE visualization from our integrated spARC diffusion operator **P_integrated_**. As done in PHATE, we first *log* normalize the diffusion operator 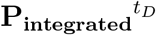 to compute potential distance **U**. In **U**, a smaller potential distance corresponds to points that are more similar to one another while higher potential distance corresponds to points that are more different as described by (18). Next, we visualize potential distances **U** into two or three dimensional coordinates via a stress-minimizing optimization procedure provided by multidimensional scaling. Using this visualization procedure, we can use spARC to visualize spatial transcripmics data more completely, potentially revealing spatial niches of distinct cell populations.

### Overview of denoising comparison

In order to compare spARC to other single cell denoising algorithms, we perform a robust set of comparisons not only visually (Fig. 1b) but also quantitatively (Fig. 2a-b). For our quantitative comparison, we utilize MERFISH which has shown to have high fidelty in capturing transcriptic counts *in situ* (31). On top of this robust dataset, we simulated increasing amounts of random Gaussian noise and drop out (Fig. 2a) and used spARC along with other algorithms to try and recapture ground truth expression dynamics from this noiseless data. For each comparison, the same alpha-decay affinity graph was utilized (adaptive bandwidth knn of 15 for expression graph and knn of 5 for spatial graph).

### Overview of signaling comparison

To establish the utility of our signaling analysis, we simulated the signaling dynamics of 634 distinct ligand-receptor interactions in our lymph node Visium data set and tried to correlate these dynamics with known downstream signaling effects (48). Comparing spARC with algorithms that either incorporate spatial location into signalling inference (SpaOTsc) or do not (CellPhoneDB), we find that spARC better correlates cytokine signaling effects with known downstream consequences of these signaling effects across ligands. While CellPhoneDB only calculates signaling interactions between cells within a cluster and is not ‘cluster-free’ like spARC and SpaOTsc, we computed single cell interaction scores by treating each barcoded spot in our Visium dataset as a cluster. After computing the interaction scores for each of the 634 interactions, we grouped interactions by ligands and computed ligand-specific interaction scores across all cognate receptors. Simultaneously, we computed ligand-specific gene signature scores by averaging the expression of all genes known to be downstream of this ligand stimulus on immune cells (48). We then computed mutual information between this ligand-specific interaction score and the ligand-specifc gene signature scores. If an interaction had high mutual information with the known down stream gene signature of that interaction, then the algorithm properly reconstructed the expected signaling dynamics.

## Code availability

The spARC package, as implemented in python, is available for download with a guided tutorial on the Krishnaswamy Lab Github page: https://github.com/KrishnaswamyLab/sparc.

## Supplementary materials

**Figure S1:**
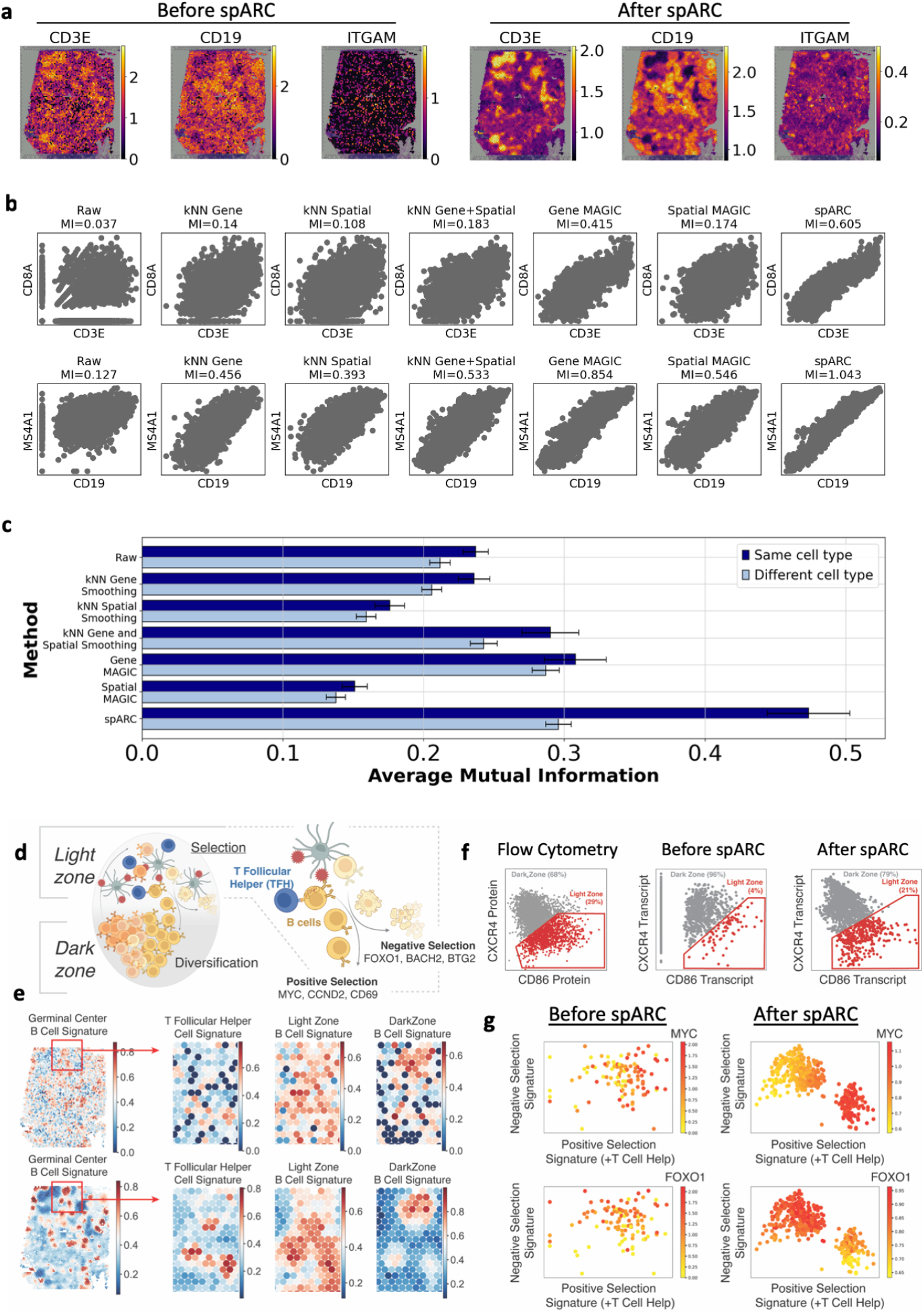
spARC recovers both shared and divergent transcriptional programs within lymph node architecture. a. spARC recovers expression of LZ (CD86) and DZ (CXCR4) markers as compared to protein expression measured by flow cytometry. b. spARC preserves divergent transcriptional programs in LZ B cells actively undergoing positive or negative TFH-mediated selection. Expression signatures were validated in (36). c. spARC recovers the expression of key celltype-specific genes from the lymph node measured with spatial transcriptomics 10X Visium platform. d. Correlating known co-expressed, cell type-specifc genes found in T cells (CD3E, CD8A) and B cells (MS4A1, CD19) after applying different denoising strategies. Mutual information between genes is used as a performance measure, indicating the degree to which gene co-expression dynamics have been recovered. e. Mutual information found between genes known to be co-expressed in the same cell type (true association) and genes known to be expressed in different cell types (false association) are measured after applying different denoising strategies. This experiment is repeated for 10 genes in each cell type, error bars indicate one standard deviation around mean mutual information for each set of comparisons. g. Schematic of germinal center architecture and postive vs. negative TFH-mediated selection made with BioRender. g. Expression of germinal center (GC) B cell, T follicular helper (TFH), light zone B cell (LZ), and dark zone B cell (DZ) expression signatures visualized in lymph node Visium tissue. LZ and DZ signatures were validated in (35).

**Figure S2:**
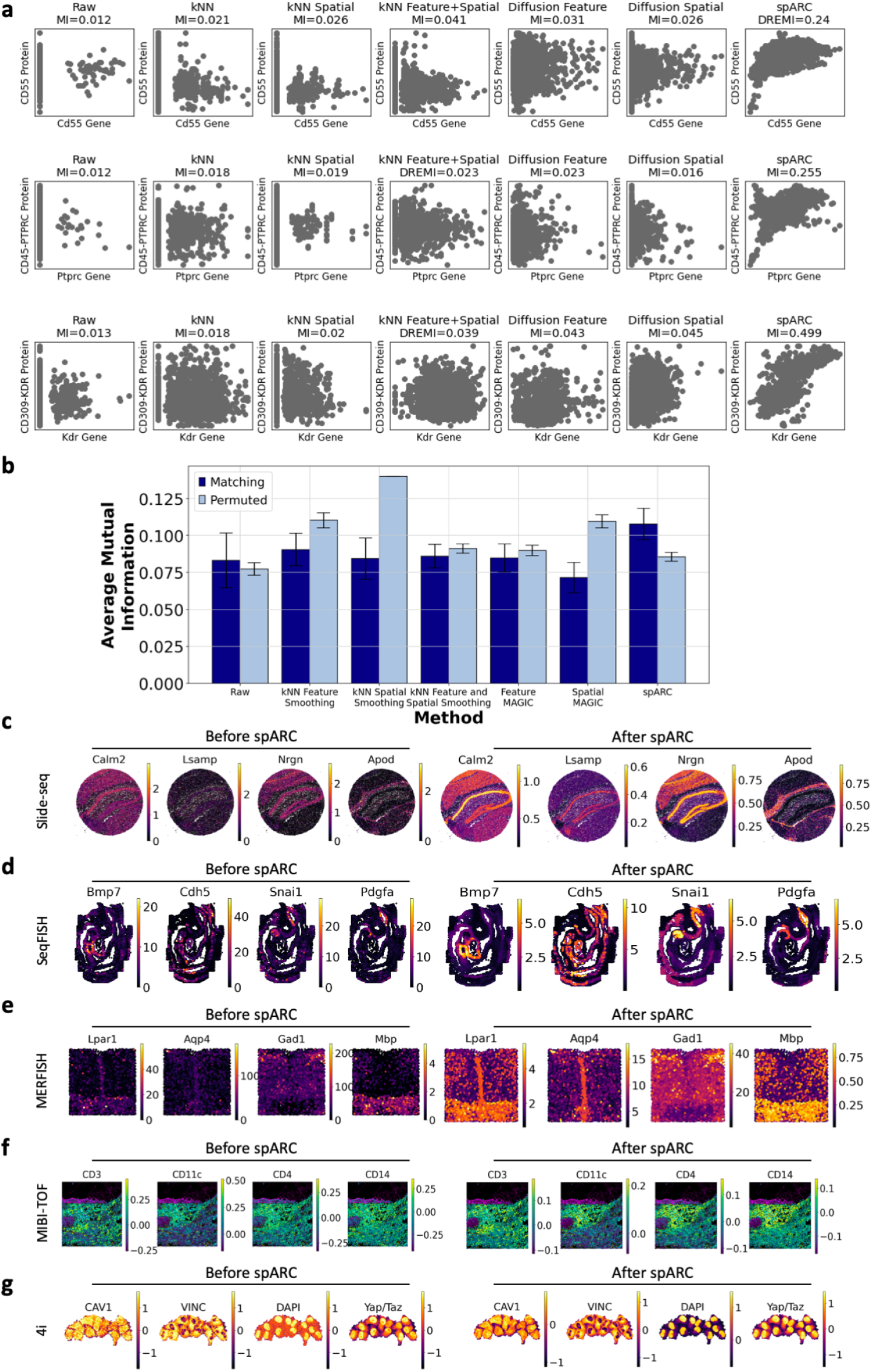
spARC recovers cross modality associations from paired gene and protein spatial data and is compatible with any spatial technology. a. Quantitative study comparing denoised results from spARC and other imputation approaches on trimodal DBIT-seq data (37) with both gene expression and protein expression measurements. Mutual information between a gene and it’s translated protein is used as a performance measure, indicating the degree to which these expression dynamics have been recovered. b. Mutual information found between genes and their translated proteins (Matching) as well as genes and different proteins (Permuted) are measured after applying different denoising strategies. Error bars indicate one standard deviation around mean mutual information for each set of comparisons. (c-g) spARC can be applied to any spatial technology including slide-seq (38), seqFISH (40), MERFISH (31), MIBI-TOF (41) and 4i (42).

**Figure S3:**
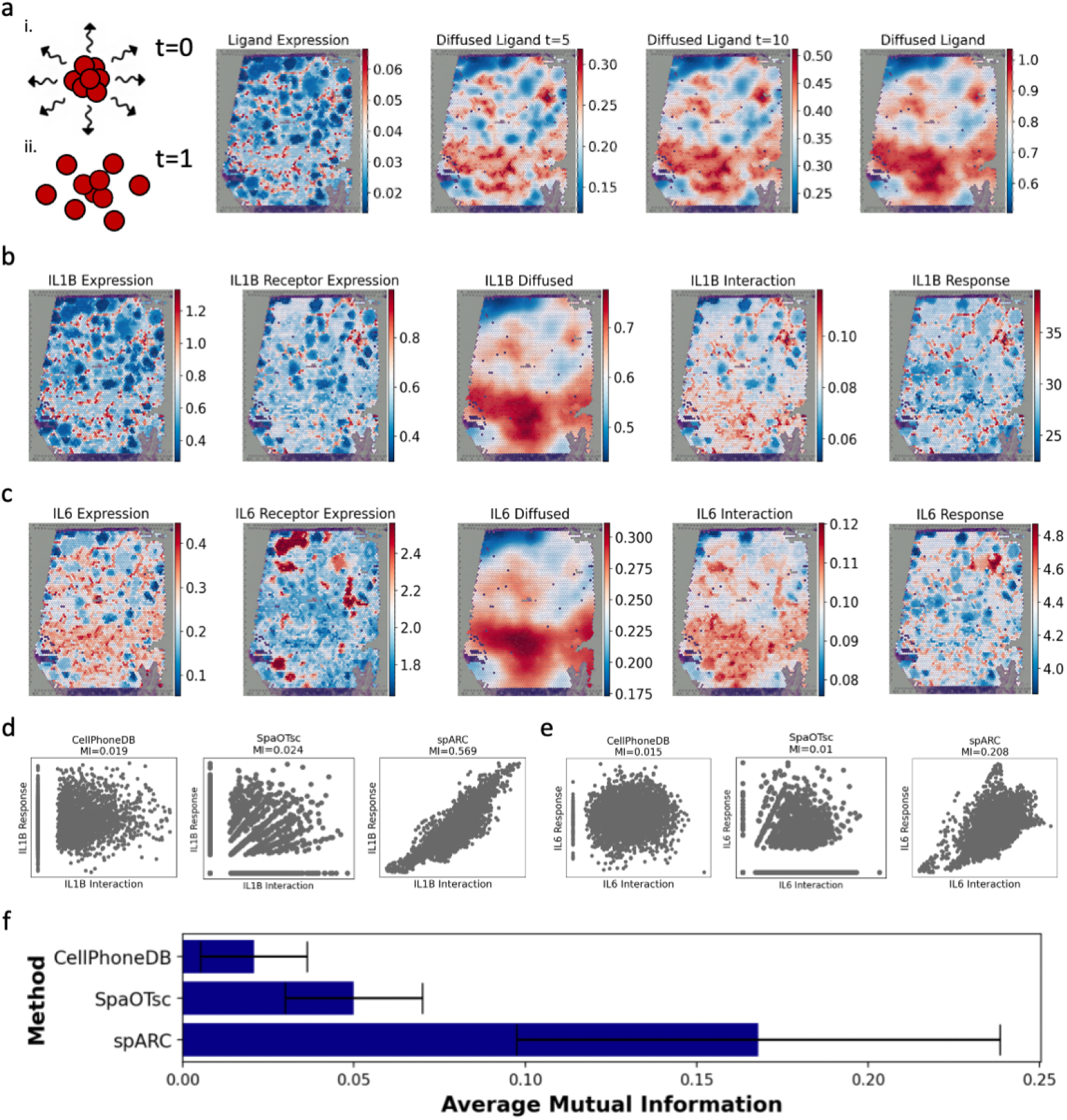
Ligand-Receptor Interactions in a Human Lymph Node. a. Illustration of diffusion of ligands in tissue as modelled by spARC across different diffusion timescales. b. Gene expression of IL1B and IL1B-receptor visualized in lymph node Visium tissue, followed by IL1B ligand diffusion in space and IL1B interaction scores as computed by spARC. Finally, IL1B response signature (see Methods) is visualized as established in (48). c. Gene expression of IL6 and IL6-receptor visualized in lymph node Visium tissue, followed by IL6 ligand diffusion in space and IL6 interaction scores as computed by spARC. Finally, IL1B response signature (see Methods) is visualized as established in (48). d. Mutual information is computed between IL1B interaction scores computed by CellPhoneDB, SpaOTsc and spARC and known IL1B signaling response signatures. e. Mutual information is computed between IL6 interaction scores computed by CellPhoneDB, SpaOTsc and spARC and known IL6 signaling response signatures. f. Mutual information between all 624 ligand-receptor interaction scores as determined by CellPhoneDB, SpaOTsc and spARC and known response signatures are computed. Error bars indicate one standard deviation around mean mutual information for each set of comparisons.

**Figure S4:**
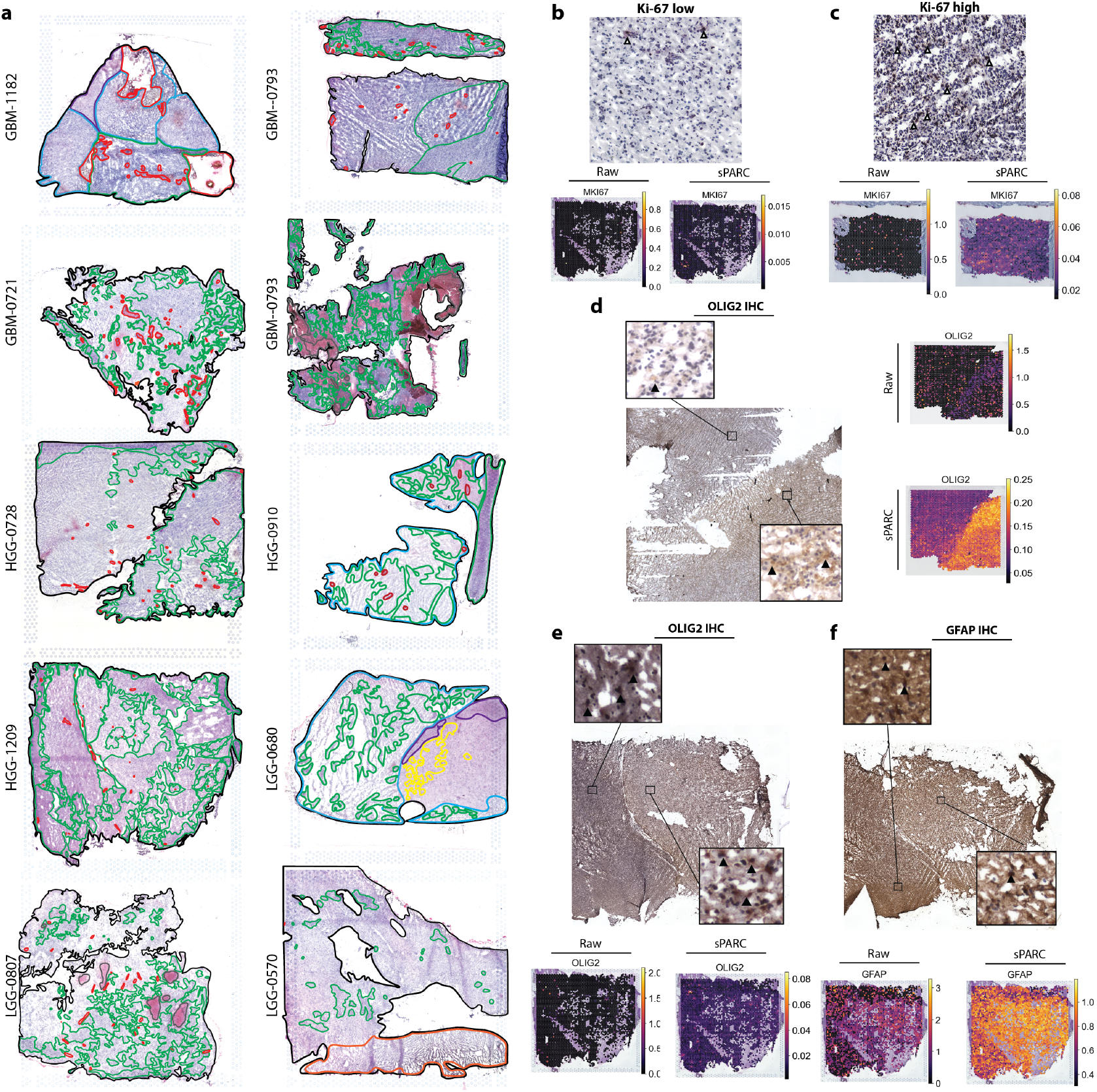
Neuropathology annotations Glioma Histology. a. All tumor histopathology is shown. In all tissue, black outlines the whole tissue, green show areas of increased cellular foci. Red demarcates areas of prominent blood vessels or microvascular profliferation. Light blue are areas of diffusily infiltrating tumor. In GB-1182, purple shows Hypercellular brain infiltrated by tumor cells. In LGG-0680, purple shows foci of high cellularity in solid tumor areas and yellow demonstrates examples of neurons surrounded by tumor. In GB-0793 and LGG-0807, grey shows zones of likely blood or other ‘leakage’ from H&E artifact c. ki-67 protein expression in HGG-1209. Arrows show cells that are positively stained for ki-67 protein. Beside this image, the raw signal of MKI67 and spARC imputed MKI67 is shown demonstrating that low expression results in non-spurious imputation of the gene in agreement with ground truth protein results. d. ki-67 protein expression in GB-0793, where white arrows are examples of positively stained cells, demonstrating uniform high expression at the protein level. The raw signal of MKI67 in this sample is sparse, but imputation with spARC restores expression to be more reflective of the ground truth. e. A whole slide image of HGG-0728 with OLIG2 protein expression. d. A whole slide image of HGG-1209 with OLIG2 protein expression uniformly. f. A whole slide image of the same sample as (d) HGG-1209 highlighting GFAP protein expression, which is more abundant than OLIG2 in this sample.

**Figure S5:**
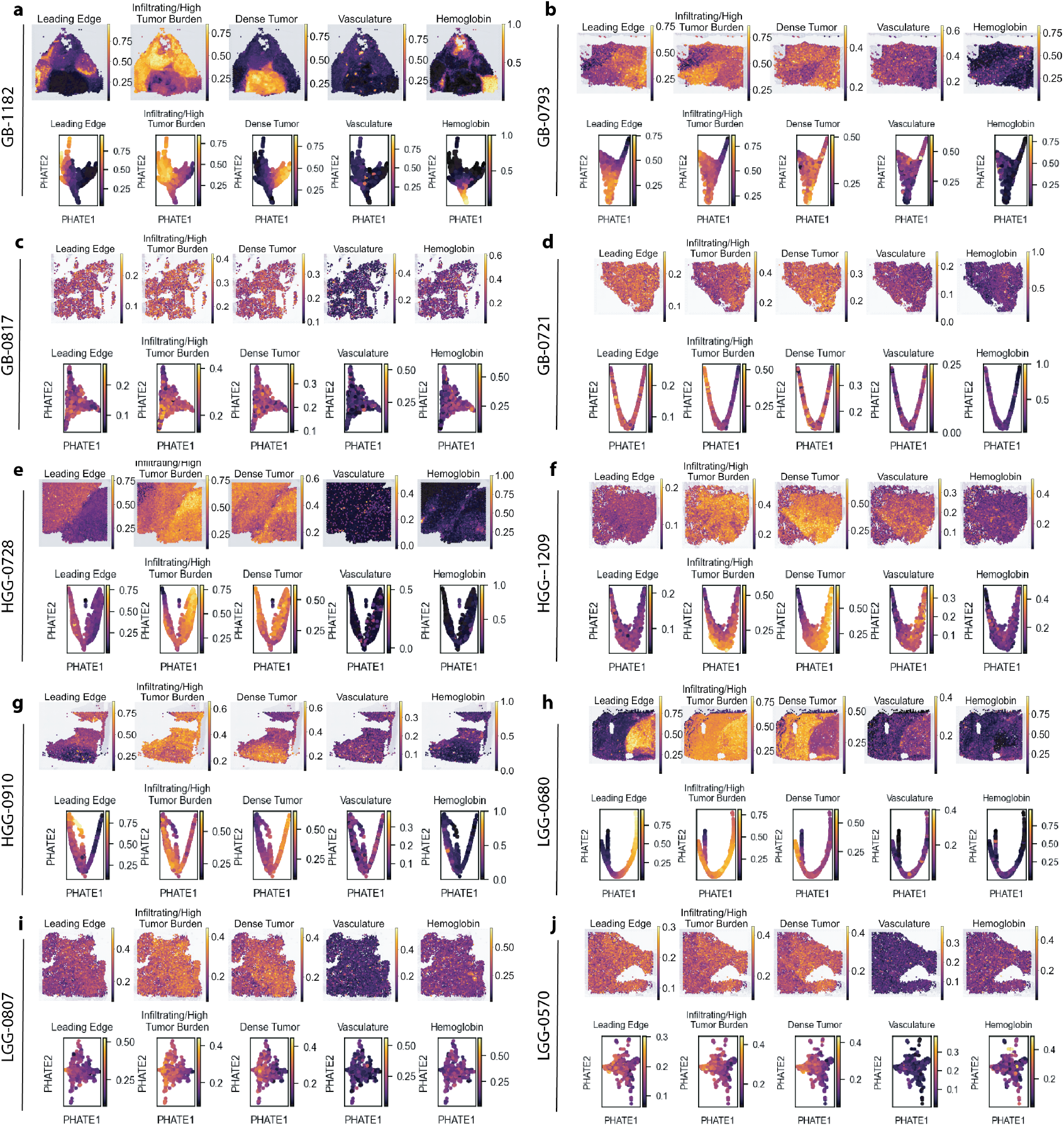
Spatial niches of IDH-MUT gliomas. Gene modules that match those used in Fig. 3 shown with composite expression on tissue (top) and using a PHATE plot (top). Both graphs are shown to demonstrate that while some samples lack spatial localization within the tissue, some gene modules show localization in the PHATE graphs, indicating the similarity between data points in low dimensional space. a. GB-1182, b. GB-0793, c. GB-0817, d.GB-0721, e. HGG-0728, f. HGG-1209, g. HGG-0910, h. LGG-0680, i. LGG-0807, j. LGG-0570.

**Figure S6:**
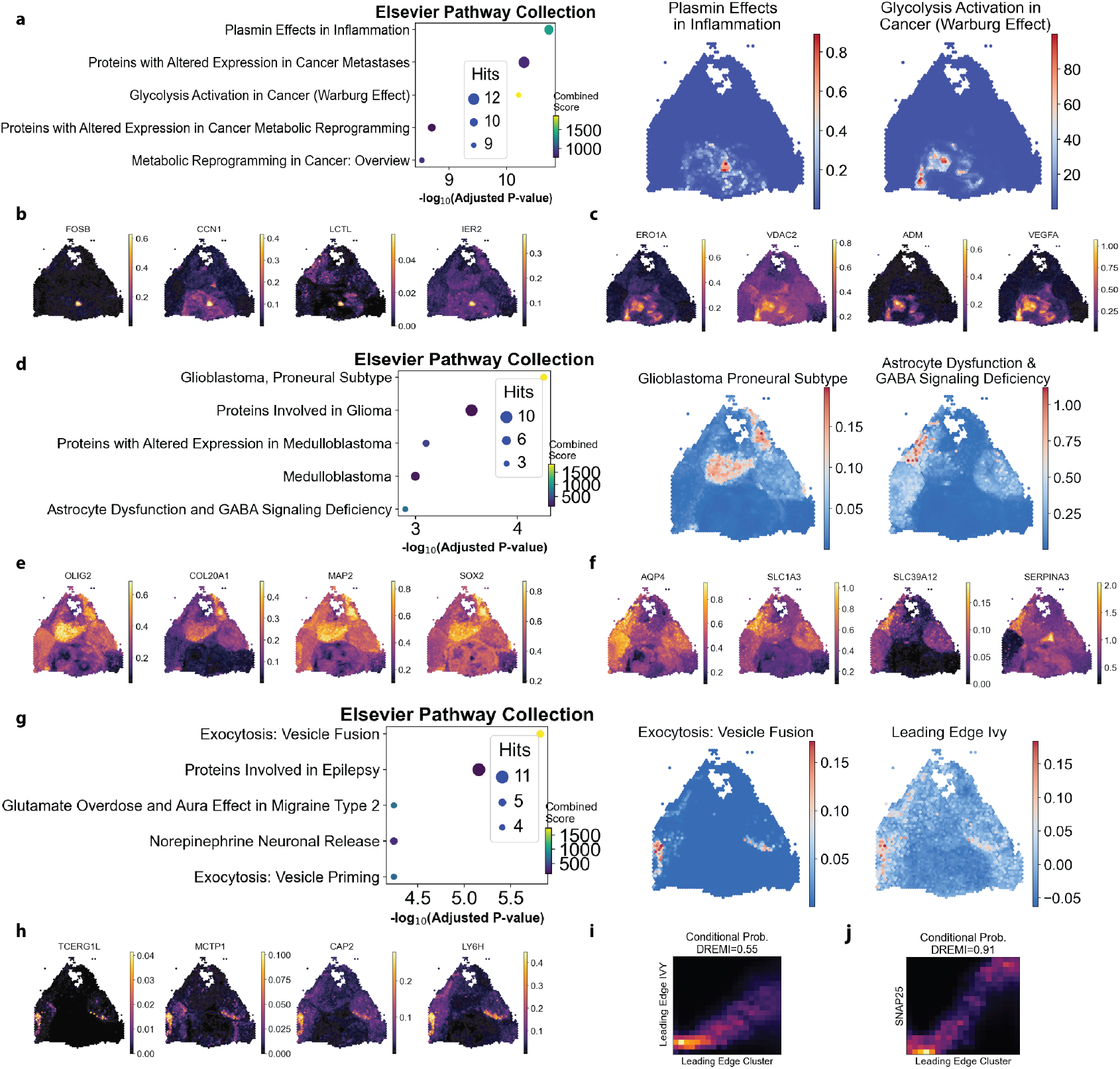
Using mutual information to identify enriched spatially localized gene modules. Left) Elsevier Pathway Collection pathway enrichment with adjusted p-value using the top 100 genes in the Right) Specific spatial expression of genes in module term. a. Dense Tumor cluster pathway enrichment b. Top genes identified using DREMI demonstrates genes with restricted expression overlapping with Plasmin Effects in Inflammation. c. Top genes identified using DREMI demonstrates genes with restricted expression overlapping with Glycolysis Activation in Cancer (Warburg Effect). d. Infiltrating/High tumor burden cluster pathway enrichment. e. Top genes identified using DREMI demonstrates genes with restricted expression overlapping with Glioblastoma Proneural Subtype module. f. Top genes identified using DREMI demonstrates genes with restricted expression overlapping with Astrocyte Dysfuction & GABA Signaling Deficiency. g. Leading Edge cluster pathway enrichment. Right image shows the median score of Leading Edge genes with a log-2 fold change of >5 and an q<0.001 (110 genes) belonging to the IVY GB database (62). h. Top genes identified using DREMI demonstrates genes with restricted expression overlapping with Exocytosis Vesicle Fusion. i. DREMI comparing the overlap between Leading Edge cluster in Leading Edge Ivy GB Database versus the top 100 genes of Leading Edge cluster in Fig. 3b. j. DREMI comparison of Leading Edge Cluster and Leading Edge marker SNAP25.

**Table S1:**
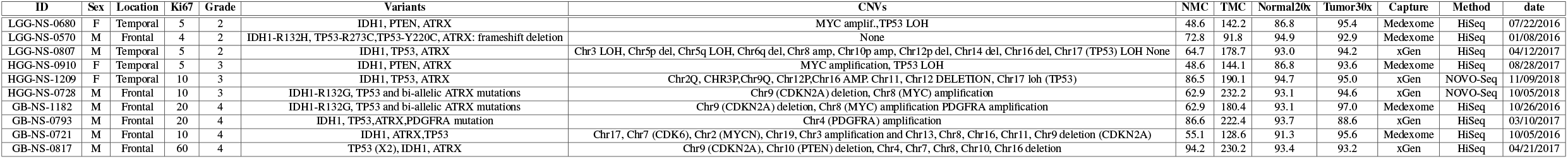
Human Glioma specimen details.

